# Antithetic integral feedback for the robust control of monostable and oscillatory biomolecular circuits

**DOI:** 10.1101/838748

**Authors:** Noah Olsman, Fulvio Forni

**Affiliations:** Department of Systems Biology, Harvard Medical School, Boston, MA 02215, USA; Department of Engineering, University of Cambridge, Cambridge CB2 1PZ, UK

**Keywords:** Synthetic Biology, Antithetic Integral Feedback, Nonlinear Control, Dominance Theory

## Abstract

Biomolecular feedback systems are now a central application area of interest within control theory. While classical control techniques provide valuable insight into the function and design of both natural and synthetic biomolecular systems, there are certain aspects of biological control that have proven difficult to analyze with traditional methods. To this end, we describe here how the recently developed tools of dominance analysis can be used to gain insight into the nonlinear behavior of the antithetic integral feedback circuit, a recently discovered control architecture which implements integral control of arbitrary biomolecular processes using a simple feedback mechanism. We show that dominance theory can predict both monostability and periodic oscillations in the circuit, depending on the corresponding parameters and architecture. We then use the theory to characterize the robustness of the asymptotic behavior of this circuit in a nonlinear setting.

## 1. INTRODUCTION

Feedback regulation is ubiquitous in biology, playing a crucial role in high-level phenomena, such as sensory perception in animals (Wiener (1948); Nakahira et al. (2019)), all the way down to the most basic processes in life, like the regulation of amino acid biosynthesis (Monod (1971)). As we refined our understanding of molecular biology, it became abundantly clear that life not only relies heavily of feedback control, but that this control is often extremely precise and robust (Barkai and Leibler (1997)). It has become a central goal of biological engineering to implement synthetic feedback controllers in cellular systems, the performance of which we hope will rival that found in nature (Del Vecchio et al. (2016)).

In the past few years, a great deal of progress has been made towards this goal of designing universal feedback controllers that can easily be used in a wide variety of contexts (Aoki et al. (2019); Samaniego and Franco (2017); Chevalier et al. (2019); Becskei and Serrano (2000); Huang et al. (2018)). Among these, a particularly promising architecture relies on what has been named the Antithetic Integral Feedback (AIF) circuit, which has been shown to implement integral feedback using a simple irreversible bimolecular interaction as the primary feedback mechanism (Briat et al. (2016)). This architecture has several appealing properties: it relies on a single molecular interaction that appears in a variety of natural contexts (e.g., the nearly irreversible sequestration of sigma factor/antisigma factor pairs), and the model of its dynamics is simple enough that it is possible to prove general results about stability and optimality (Aoki et al. (2019); Olsman et al. (2019a)). Recent theoretical work has focused both on global results for arbitrary plant-controller systems and local results focused on applying classic tools from control theory to simplified models of closed-loop dynamics (Qian et al. (2018); Qian and Del Vecchio (2018); Olsman et al. (2019a,b); Briat et al. (2016, 2018)).

While much progress has been made, there are still some basic properties of the system that have, so far, proven difficult to address theoretically. For example, it was shown via simulation that, for some plant models, local instability around a unique equilibrium point of the closed-loop system yields a single stable limit cycle globally (Briat et al. (2016)). While we have a fairly good understanding of local stability for some biologically realistic parameter regimes, there is still relatively little that is understood about the specifics of the AIF circuit’s global, nonlinear behavior. This is in part due to the fact that the simplest models of the system that can produce limit cycles have at least four states (Olsman et al. (2019a)), making it difficult to use any classical results from the theory of planar systems (e.g., the Poincare-Bendixson theorem). Notably, the recent paper (Margaliot and Sontag (2019)) proved the existence of compact attractors using a generalized version of Poincaré-Bendixson for cooperative systems with respect to high-rank cones. Their approach is based on the sign pattern of the system Jacobian, however it is unclear how to extend it to model classes that do not satisfy this sign pattern. It has been similarly difficult to construct a global Lyapunov function that would facilitate the use of other nonlinear tools, such as LaSalle’s Invariance Principle or a generalized Hopf bifurcation analysis.

To address these issues, we will use the tools of dominance theory to study the AIF circuit. Dominance analysis is closely related to contraction methods and those based on invariant cone methods (e.g., Smith (1979, 1980); Mallet-Paret and Smith (1990); Wang and Slotine (2005); Sanchez (2009); Lian and Wang (2015)). For an extensive literature review and a detailed discussion of the relationship between these methods and dominance analysis, see Forni and Sepulchre (2019). By adopting a differential perspective, dominance theory generalizes classical tools from linear system theory to the nonlinear setting, for the analysis of multistable and oscillatory closed-loop systems. In this paper we will illustrate how to use dominance theory to derive novel results about a biological system that, so far, has proven difficult to analyze with classical tools. We show how the AIF system achieves closed-loop regulation and how the same device can be used to design robust oscillatory circuits. The analysis emulates the approach pursued for mechanical and electrical system (Forni and Sepulchre (2014); Miranda-Villatoro et al. (2018b)), demonstrating the potential dominance theory has for robustness analysis of AIF circuits. While our results focus on a simple 2-state plant model (with both a linear and non-linear variant) for pedagogical reasons, it should be noted that these results generalize quite naturally to plants of arbitrary dimension and with any smooth class of nonlinearities.

## 2. ANTITHETIC INTEGRAL FEEDBACK CIRCUITS

The AIF controller, which we denote Σ_*z*_, follows the dynamics:

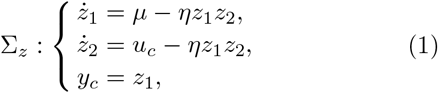

where *z*_1_ ≥ 0 can be thought of as the actuator species and *z*_2_ ≥ 0 the sensor species. *μ* is the reference input (for adaptation). *η* is the rate at which *z*_1_ and *z*_2_ bind and are jointly removed from the system. *η* typically captures a sequestration rate, but can also represent any interaction in which two species are mutually inactivated. The key property here is that *η* is exactly identical for both *z*_1_ and *z*_2_.

Neither *z*_1_ nor *z*_2_ in system (1) directly implements an integrator. Their difference *z* = *z*_1_ − *z*_2_, however, follows the dynamics

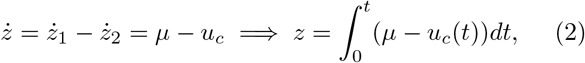

which encodes a virtual integrator internal to the system’s state. At equilibrium, we have that

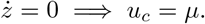

This means that, for any plant 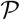 with input *u* and output *y*, any stable closed-loop interconnection given by

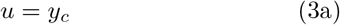

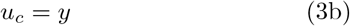

must asymptotically yield *y = μ*, demonstrating the robustness of the closed-loop equilibrium to parametric variations.

For the purposes of this article, we will study the Jacobian of system (1):

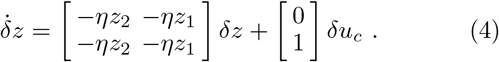

A crucial observation is that system (4) is not the linearization of the system around a specific equilibrium. Rather, it is the linearization of the system along any possible trajectory *z*(·) of system (1). The analysis in the next sections depends crucially on this general parameterization of the Jacobian (i.e., a differential perspective of dynamics).

## 3. DOMINANCE THEORY

### 3.1 The workflow of dominance analysis

Dominance theory provides a set of tools to study non-linear models which have behavior that is not constrained to a single stable equilibrium Forni and Sepulchre (2019); Miranda-Villatoro et al. (2018a). The theory takes non-linear models with parameters in a certain range and characterizes their behavior through Lyapunov-like linear matrix inequalities (LMIs). The intuition for the theoretical framework is that the dynamics of a *p*-dominant system

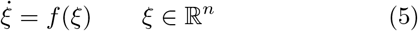

can be split into fading sub-dynamics of dimension *n* − *p* and dominant sub-dynamics of dimension *p*, the latter constraining the system’s steady-state behavior. The attractors of a *p*-dominant system must be compatible with a system of dimension *p*. For 1-dominant systems this means that every bounded trajectory converges to an equilibrium. For 2-dominant systems, bounded trajectories converge to a simple attractor, compatible with a planar dynamics.

Dominance is studied differentially, by looking at the linearized dynamics along system trajectories, namely

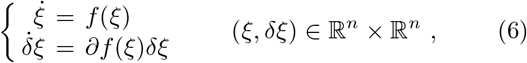

where *∂f*(*ξ*) is the Jacobian of *f* at *ξ*^1^.

#### Definition 1.

The nonlinear system (5) is *p*-dominant with rate λ ≥ 0 if there exist a symmetric matrix *P* with inertia (*p*, 0, *n* − *p*) and a positive constant *ε* such that

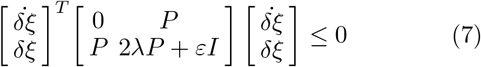

along the trajectories of (6).

We recall that a matrix 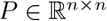 with inertia (*p*, 0, *n* − *p*) has *p* negative eigenvalues and *n* − *p* positive eigenvalues. (7) is equivalent to finding a uniform solution *P* to the matrix inequality

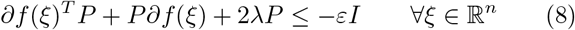

(for some *ε* > 0). Inequality (8) is a standard Lyapunov inequality adapted to the linearization, where positivity of *P* has been replaced by a constraint on the matrix inertia. The constraint on the inertia is used to separate dominant and fading dynamics. In fact, for linear systems *f*(*ξ*) = *Aξ*, LMI (8) implies that *A* has *p* dominant eigenvalues to the right of −λ, and *n* − *p* stable eigenvalues to the left of −λ, (Forni and Sepulchre, 2017, Proposition 1).

The use of dominance theory in nonlinear control is motivated by the following theorem

#### Theorem 1.

(Forni and Sepulchre, 2019, Corollary 1).

Every bounded solution of a *p*-dominant system with rate λ ≥ 0 asymptotically converges to

- a unique fixed point if *p* = 0;
- a fixed point if *p* = 1;
- a simple attractor if *p* = 2, that is, a fixed point, a set of fixed points and connecting arcs, or a limit cycle.

It is important to note that LMI (8) is numerically tractable. As shown in Boyd et al. (1994), it can be reduced to a finite set of inequalities built from a finite set of linear matrices {*A*_1_, …, *A_N_*} that satisfy the property

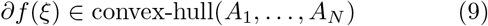

for all 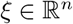. For the dynamics considered in this paper, these matrices *A_i_* can be easily obtained from the system Jacobian *∂f*(*ξ*) computed at a set of points forming a convex hull of the region of interest (both in parameter space and state space). The whole workflow is illustrated in Figure 1.

**Fig. 1.**
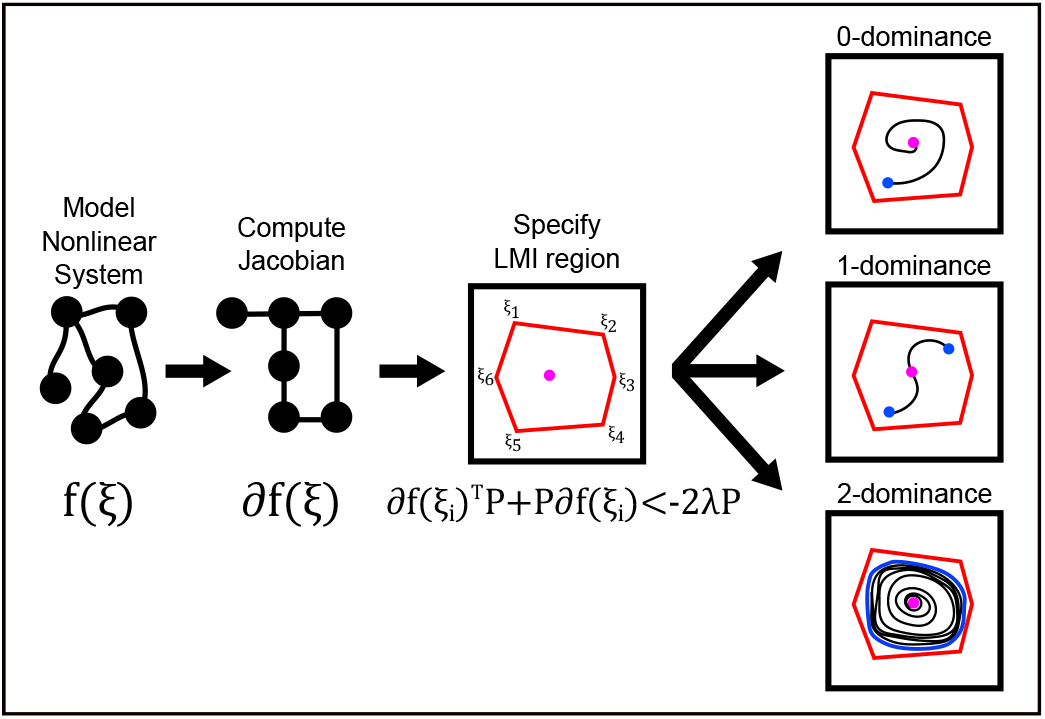
The workflow of dominance theory: 1) start with a nonlinear model of interest; 2) compute the corresponding Jacobian; 3) compute the convex hull matrices {*∂f*(*ξ*_1_), …,*∂f*(*ξ_N_*)}. For the dynamics considered in this paper this reduces to specify points *ξ_i_*, the convex hull of which define a region of interests (red); finally 4) use LMIs to generate a certificate of dominance that describes the system’s limiting behavior within the region (blue). In the last panel, we see that 0-dominance corresponds to single equilibrium, 1-dominance corresponds to multiple equilibria, and 2-dominance corresponds to a simple attractor.

### 3.2 Dominance design through necessary conditions

The eigenvalues of the Jacobian *∂f*(*x*) of a *p*-dominant system are always split into two groups, with *p* eigenvalues to the right of −λ and the remaining ones to the left (at each *x*). This necessary condition can be studied via classical control tools like root locus method and Nyquist criterion, which thus provide relevant insights for parameter tuning of AIF-based controllers.

Consider the (open) nonlinear system

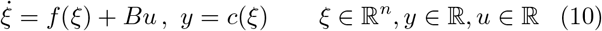

and the closed-loop system arising from (10) through the feedback interconnection

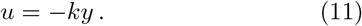

If the closed-loop system (10),(11) is *p*-dominant with rate λ then the closed-loop Jacobian

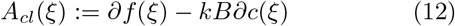

satisfies (8) and the splitting of its eigenvalues can be predicted through root locus analysis and Nyquist criterion applied to the (frozen) state-dependent transfer function

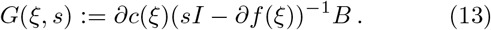

#### Theorem 2.

(Root locus and Nyquist criterion).

Suppose that for some *k* = *k*\* the closed-loop system described by (10),(11) is *p*-dominant with rate λ. Suppose that the pairs (*∂f*(*ξ*), *B*) and (*∂f*(*ξ*), *∂c*(*ξ*)) are controllable and observable at *ξ*, respectively. Then,

- the root locus of *G*(*ξ, s*) for the feedback gain *k*\* has *p* roots whose real part is greater than −λ and *n* − *p* roots whose real part is smaller than λ;
- the Nyquist plot of *G*(*ξ*, −λ + *jω*) for 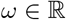 encircles the point 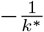 exactly (*p* − *q_ξ_*) times in the clockwise direction, where *q_ξ_* is the number of poles of *G*(*ξ, s*) whose real part is greater than −λ.

**Proof**. Under controllability and observability assumptions, the first item follows directly from the fact that the roots of 1 + *kG*(*ξ, s*) correspond to the eigenvalues of *A_cl_*(*ξ*). The second item follows from the argument of the proof of (Miranda-Villatoro et al., 2018a, Theorem 3.1).

In what follows we will use the necessary conditions of Theorem 2 as guidelines for control design, since they define minimal requirements that a dominant closed-loop system has to satisfy. Root locus and Nyquist diagrams will be used to find model parameters that guarantee a suitable splitting of the closed-loop eigenvalues, taking into account classical robustness considerations. Validation will then be certified through linear matrix inequalities (8) applied to the closed-loop Jacobian in equation (12).

## 4. FIRST-ORDER PRODUCTION SYSTEM: HOMEOSTASIS AND OSCILLATIONS

### 4.1 Linear feedback

We will start by analyzing the linear plant model

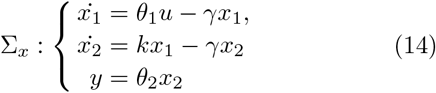

We refer to (14) as *first-order production* system. Here *θ*_1_ and *k* represent first-order production rates for each species, and *γ* represents a common first-order degradation rate. It has been demonstrated via simulation that this simple model in closed loop with the nonlinear AIF controller can exhibit stable limit cycles (Briat et al. (2016)), which appear to arise when the linearized closed-loop system becomes locally unstable. In the limit of large *η* this corresponds to the parametric condition 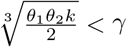, as shown in (Olsman et al. (2019a)). We will then extend our study to a plant with the same qualitative architecture but where the interconnections are modeled with more realistic saturation behavior, such as with the Hill function *θ*_1_(*u*) = *u^N^*/(*k*_1_ + *k*_2_*u^N^*).

We represent the closed loop of AIF circuit and first-order production system (1), (3), (14) as the interconnection of (1), (3a), (14) with input 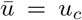 and output 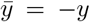, through negative feedback 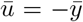. The linearized closed loop is shown in Figure 2. The state-dependent linearized transfer function from 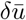 to 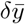 reads

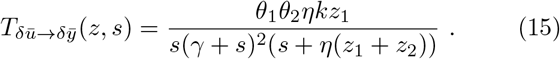

**Fig. 2.**
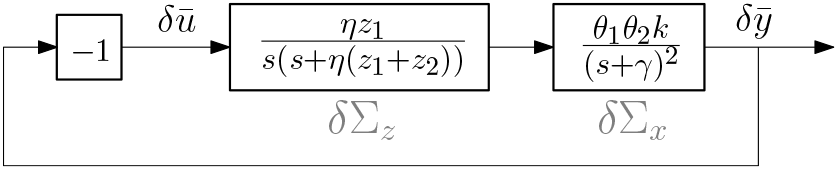
Linearized closed-loop system diagram of (1), (3a), (14) with 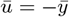.

For any fixed *z*, the root locus of 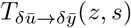 shows that poles move towards infinity as *θ*_1_*θ*_2_*k* becomes large, along four asymptotes oriented as 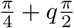 where *q* ∈ {0, 1, 2, 3}. In agreement with Theorem 2, this suggests a potential transition from 0-dominance to 2-dominance, as *θ*_1_*θ*_2_*k* increases, with poles separated into stable and dominant pairs, respectively to the left and to the right of the (*z*-dependent) centroid. Similar observations follow from the Nyquist plot, which makes no rotations around −1 for *θ*_1_*θ*_2_*k* small, but has two clockwise rotations around −1 for large *θ*_1_*θ*_2_*k*.

For simplicity, we develop the details of the analysis for fixed parameters *μ* = 2, *η* = 10 (controller) and *θ*_1_ = 1, *γ* = 1 (plant). The analysis can be easily adapted to different parameters.

As a first case, consider *θ*_1_*θ*_2_*k* = 1 (small). The Nyquist plots in Figure 3(d) samples 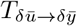 for *z* constrained to the convex red region 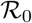 in Figure 3(b). Indeed, accordingly to Theorem 2, the Nyquist plots are all compatible with 0-dominance. Compatibility is reinforced by Figure 3(c), which shows that the closed-loop Jacobian eigenvalues have negative real part for all 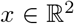 and all 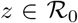. From the perspective of the LMI (8), the closed loop is 0-dominant with rate λ = 0 for 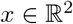 and 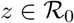. This is certified by the first positive-definite matrix *P* in Table 1, which is a solution to (8) for *ξ* = [*z^T^,x^T^*]^*T*^ and for *f* representing the closed-loop dynamics. The solution is computed by convex relaxation, taking advantage of the finite number of vertices of 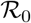.

**Fig. 3.**
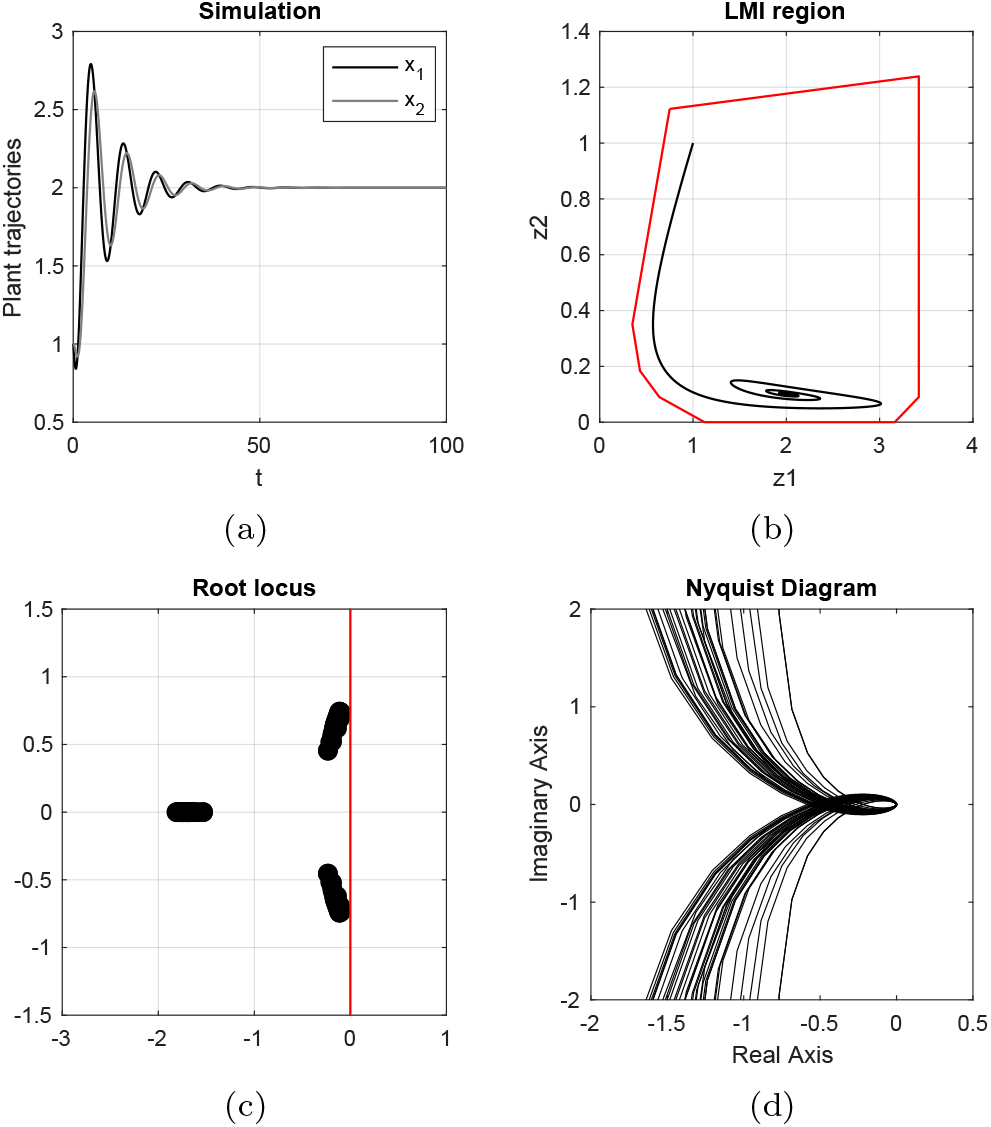
(a,b): Trajectories (time-trajectories for *x*, *z*-plane projection) for *θ*_2_ = *k* = 1. The red curve in (b) delimits the boundary of the convex region 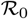. (c): closed-loop Jacobian eigenvalues for 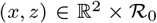. (d): Nyquist locus of 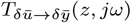 for 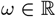 and 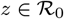.

**Table 1.** Matrices *P* for dominance, for different values of parameters *θ*_2_ and *k*.

|  |  |
| --- | --- |
| $\theta_2 = 1$ | $\begin{bmatrix} 82.7650 & -79.6308 & -24.2526 & -42.5767 \\ -79.6308 & 89.5784 & 24.7259 & 43.0863 \\ -24.2526 & 24.7259 & 79.5742 & 24.4801 \\ -42.5767 & 43.0863 & 24.4801 & 83.9389 \end{bmatrix}$ |
| $k = 1$ | |
| 0-dom |  |
| $\theta_2 = 4$ | $\begin{bmatrix} -6.0678 & 18.1011 & -60.7826 & 57.3881 \\ 18.1011 & -14.2661 & 60.3814 & -57.2023 \\ -60.7826 & 60.3814 & 0.7924 & -42.0355 \\ 57.3881 & -57.2023 & -42.0355 & -228.9326 \end{bmatrix}$ |
| $k = 1$ | |
| 2-dom |  |
| $\theta_2 = 1$ | $\begin{bmatrix} -15.2507 & 32.0157 & -83.0632 & 26.8417 \\ 32.0157 & -26.7504 & 82.5277 & -26.8390 \\ -83.0632 & 82.5277 & -4.1886 & -15.9095 \\ 26.8417 & -26.8390 & -15.9095 & -26.8531 \end{bmatrix}$ |
| $k = 4$ | |
| 2-dom |  |

From Theorem 1, it follows that every attractor in 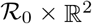 is necessarily a fixed point, as illustrated by plant time trajectories in Figure 3(a) and controller phase plane trajectories in Figure 3(b). Local asymptotic stability of the equilibrium 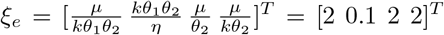 follows from standard Lyapunov argument, using (*ξ − ξ_e_*)^*T*^*P*(*ξ − ξ_e_*) as a Lyapunov function. We observe that the region admits a unique fixed point, since the presence of two fixed points *ξ_e_* and 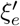 would mean

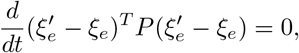

which is not compatible with LMI (8).

Figure 3(d) makes clear that 0-dominance is lost by increasing the feedback gain *θ*_1_*θ*_2_*k*. Consider the convex red region 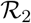 defined by the red curve in Figure 4(b) and take *θ*_1_*θ*_2_*k* = 4. For 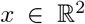 and 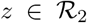, Figure 4(e) shows that the closed-loop Jacobian eigenvalues are not compatible with 0-dominance but still support 2-dominance with rate λ = 1 (from Theorem 2). This is reinforced by the Nyquist plots in Figure 4(f), which satisfies Theorem 2. From the perspective of the LMI (8), the closed-loop system is 2-dominant with rate λ = 1 for 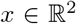 and 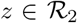, as certified by the second and third matrices in Table 1.

**Fig. 4.**
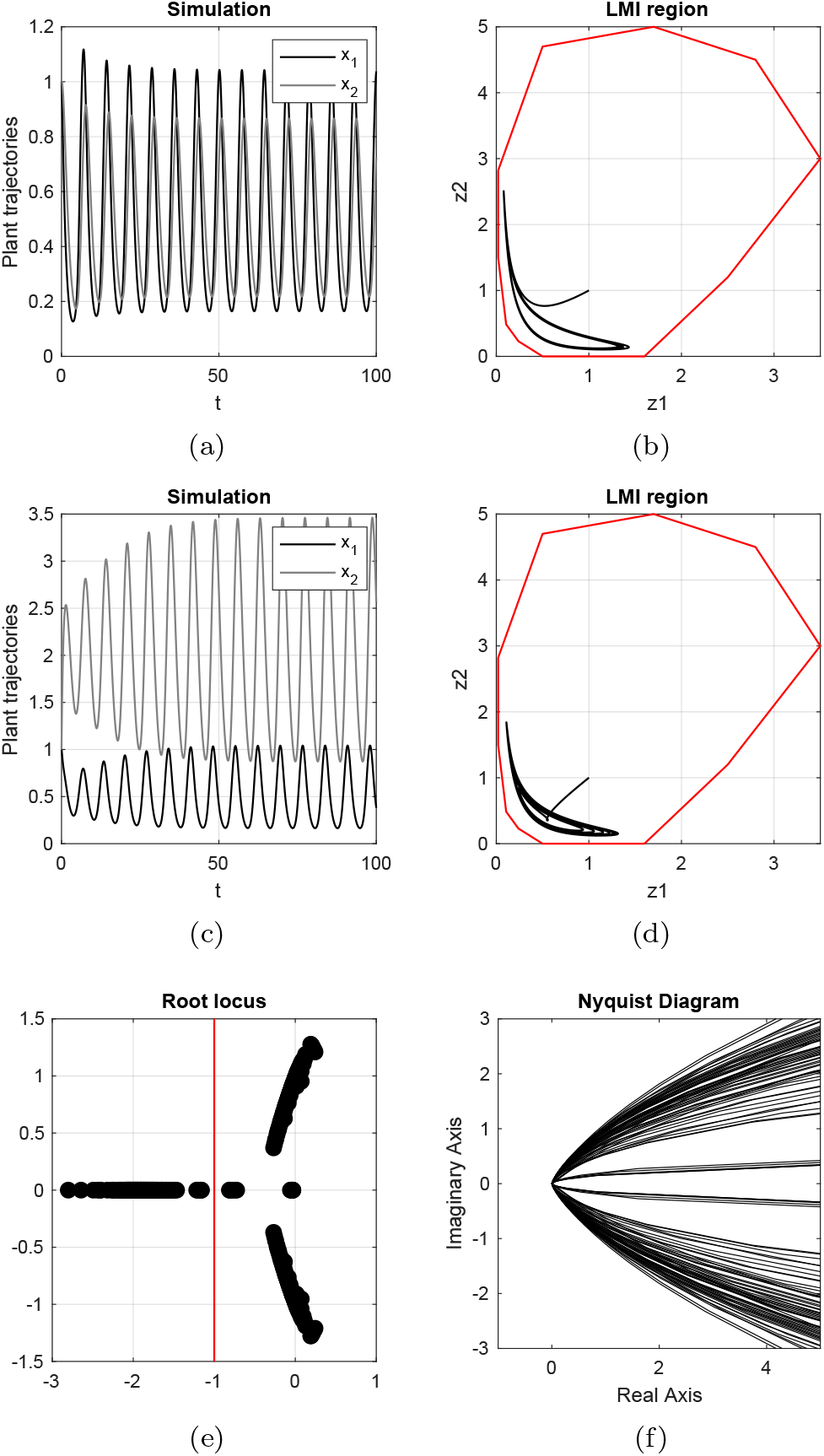
(a,b): Trajectories for *k* = 1,*θ*_2_ = 4. (c,d): trajectories for *θ_2_* = 1, *k* = 4. The red curve in (b,d) delimits the convex region 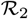. (e): closed-loop Jacobian eigenvalues for 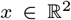 and 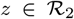. (f): Nyquist locus of 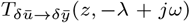, for 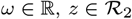, and λ = 1.

From Theorem 1, any attractor in 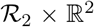 is a simple attractor. Furthermore, since the region contains only unstable equilibria (by computing the closed-loop Jacobian eigenvalues for 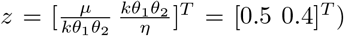, the attractor must be a limit cycle, as illustrated by Figures 4(a)–4(d).

The analysis above shows how the feedback gain (of the linearization) modulates between monostable and oscillatory behaviors. Gain tuning is based on intuitive linear feedback considerations, from Nyquist diagrams and root locus argument, followed by formal certification through linear matrix inequalities. The analysis illustrates the versatility of the antithetic integral control, which enables circuits capable of homeostatic regulation but also of stable oscillations.

### 4.2 Nonlinear feedback and robustness

The simplest way to develop robustness analysis is to encode perturbations via linear matrix inequalities (8). This can be achieved by considering a perturbed closed-loop Jacobian *∂f*(*ξ*) + Δ(*ξ*) where Δ captures a set of unknown bounded perturbations 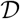 on the system Jacobian.

The existence of a uniform solution *P* to

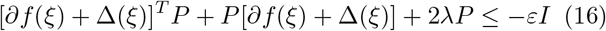

with fixed inertia for any *ξ* any 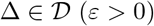 guarantees robust dominance. As in LMI (8), such a *P* can be found through convex relaxation, based on the linearization of the system computed at a finite number of points, both in state space and in parameter space.

In general, several classical tools from robust linear control have been extended to dominant systems, like robustness margins, circle criteria, and small gain theorem. The intuition is provided by the continuity of Nyquist diagrams and root locus with respect to perturbations, which shows how small perturbations do not change the dominance of the system (as in classical linear robust analysis). The interested reader is referred to Padoan et al. (2019a,b); Miranda-Villatoro et al. (2018a); Forni and Sepulchre (2019).

For the plant in Section 4.1, we study the closed-loop behavior in the presence of nonlinearities and uncertainties by considering (nonlinear) perturbations on the sequestration rate *η* and by replacing the linear feedback *θ*_1_(*u*) = *θ*_1_*u* with a nonlinear saturation such as the Hill function

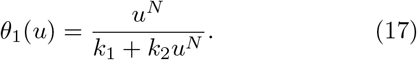

From (17), the state-dependent open loop transfer function reads

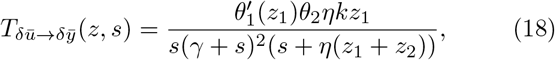

which shows that the saturation essentially modulates the (linearized) feedback gain. *θ*_1_ is non-decreasing thus, from Theorem 2, the Nyquist plots in Figure 3(d) remains compatible with 0-dominance whenever 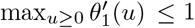. Likewise, Figure 4(f), shows that compatibility with 2-dominance should be guaranteed for any 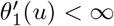.

A complete analysis is beyond the scope of the current paper. We look only at the two specific Hill functions in Figures 5(a) and 5(b). The left one satisfies 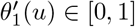, thus it is compatible with 0-dominance. The right one has a peak 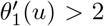, which is compatible with 2-dominance but not with 0-dominance.

**Fig. 5.**
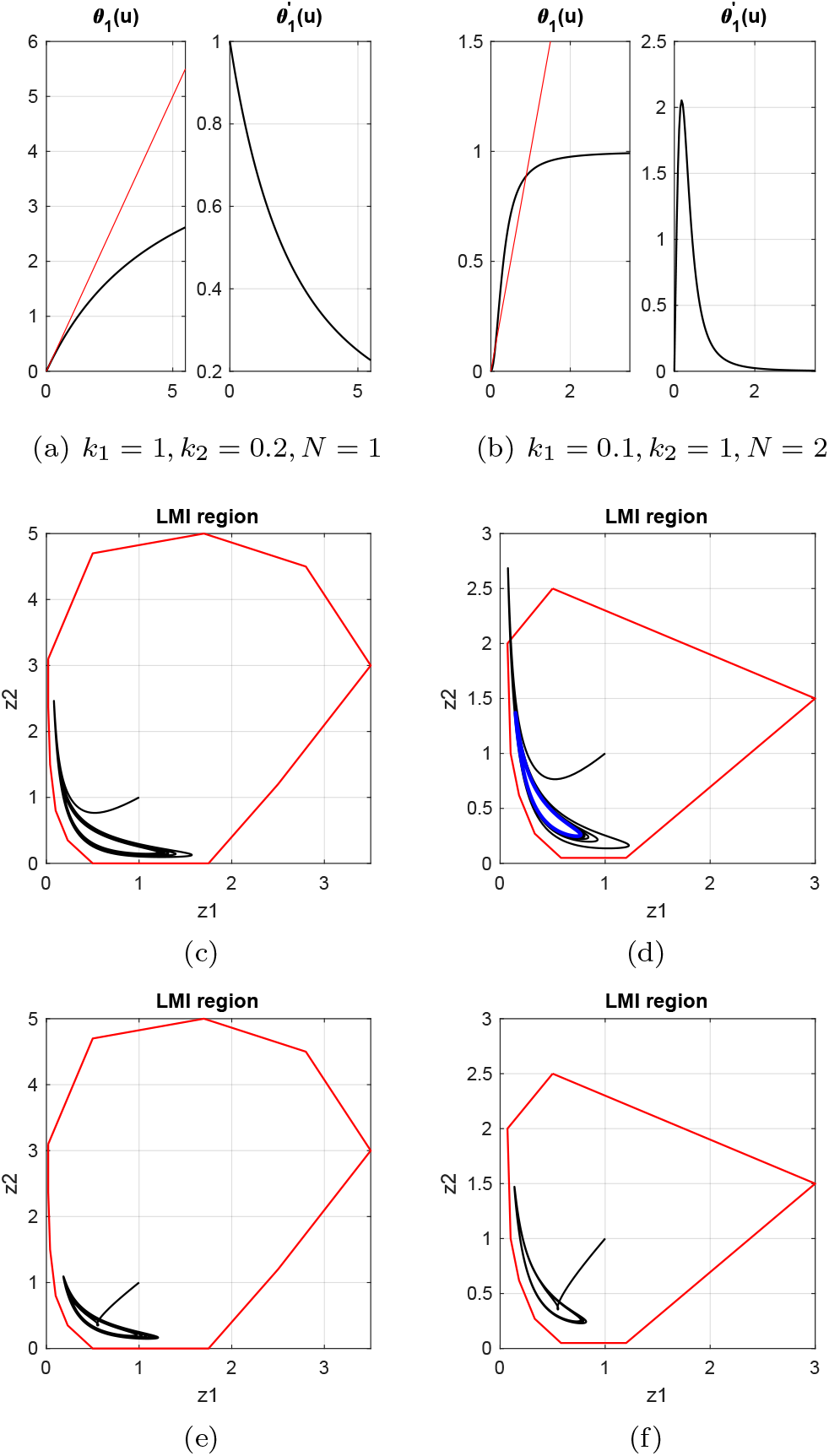
(a,b) - Hill functions. (c,d,e,g) - Trajectories projected on the *z*-plane. 2-dominance holds for *z* constrained within the red curve and *x* unconstrained. Left column: simulations related to Hill function (a). Right column: simulations related to Hill function (b). (c,d) refer to *θ*_2_ = 4, *k* = 1. (e,f) refer to *θ*_2_ = 1, *k* = 4. The blue curve in (d) shows the location of the limit cycle projected on the *z* axes, for clarity.

The LMI (16) certifies the intuitive argument from the transfer function. For the Hill function in Figure 5(a) and for *θ*_2_ = *k* = 1, 0-dominance is preserved in a sizeable region around the (new) fixed point. Likewise, 2-dominance is preserved for both Hill functions, as shown in Figure 5. The figure shows how the nonlinearities affect shape and position of the attractor but also the region of feasibility of the LMI, as illustrated by the reduced red region in Figures 5(d) and 5(f).

To complete the analysis, we briefly look into robustness to parametric uncertainties. For reasons of space we consider only the closed loop based on the Hill function in Figure 5(a). We replace the term *ηz*_1_*z*_2_ in (1) with the uncertain monotonic nonlinear binding

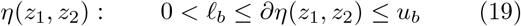

From the transfer function (18) it is clear that the Nyquist diagram does not change significantly. Indeed, for ℓ*_b_* = 7 and *u_b_* = 13 (30% variation on the nominal value), the LMI (16) certifies 2-dominance for *z* constrained within the red region of Figure 6.

**Fig. 6.**
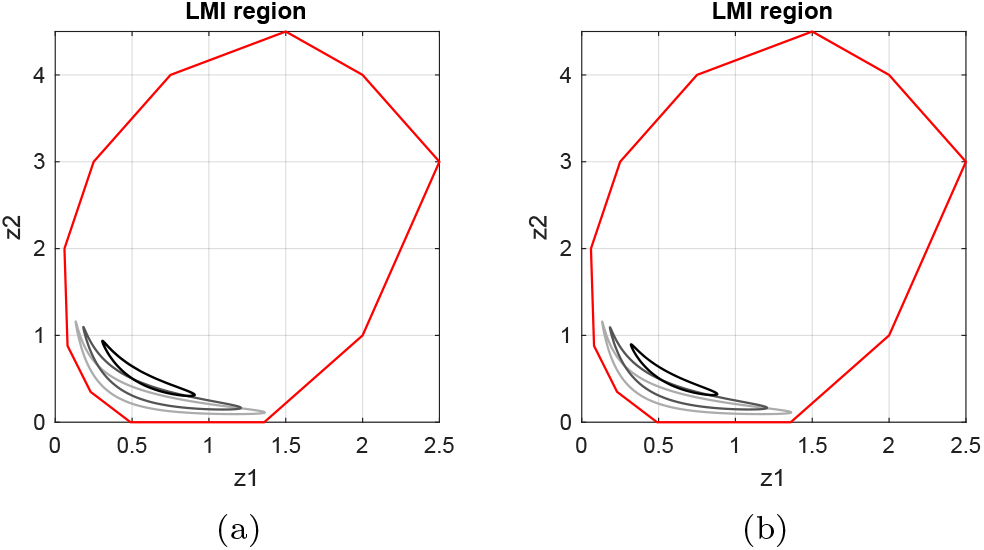
(a) *θ*_2_ = 4, *k* = 1; (b) *θ*_2_ = 1, *k* = 4. For simplicity, simulations use *η*(*z*_1_*z*_2_) = *ηz*_1_*z*_2_ for different values *η* ∈ {7, 10, 13}. The shaded gray attractors correspond to these three values, for the Hill function activation in Figure 5(a). 2-dominance holds for *z* constrained within the red curve and *x* unconstrained.

We remark that achieving robust 2-dominance does not guarantee that oscillations persist, since the attractor may reduce to an equilibrium. A sufficient condition for the attractors within the region of 2-dominance to be limit cycles is that every equilibrium in the region is unstable. This holds for the two Hill functions of Figure 5.

## 5. FUTURE DIRECTIONS

There are a number of natural directions to extend this work. For the sake of clarity we focused so far on a constrained model class, developing a detailed analysis of a few example cases. However, there is no fundamental barrier to analyzing other biological systems with similar control structure. For example, Chen and Arkin (2012) showed that it is possible to use an AIF architecture along with positive feedback to produce bistability, such as in the model

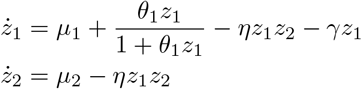

Where *μ*_1_, *μ*_2_ are transient inputs that can be used to switch the system from one state to the other. It is likely the case that such an architecture can be shown to exhibit 1-dominance, and that even more complex plant models with the same positive feedback will still reduce to 1-dominant dynamics.

Alternatively, in Olsman et al. (2019a) a plant described by the dynamics

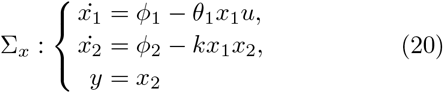

was shown via simulation to exhibit both a single locally stable equilibrium and a stable limit cycle for a particular set of parameters. It is likely possible to demonstrate this unusual form of bistability via dominance analysis. We would expect that such a system should not be described by global dominant behavior, but that it is possible to construct regions of 0- and 2-dominance that would characterize the behavior of the system within those specific regions. Indeed, simulations show that the closed loop (1), (3), (20) has stable oscillations for *μ* = 2, *η* = 10, *θ*_1_ = *k* = *ϕ*_1_ = *ϕ*_2_ = 1, and *θ*_2_ = 4, and our preliminary analysis shows that the closed loop is 2-dominant in the neighborhoods of its limit cycle, as illustrated in Figure 7.

**Fig. 7.**
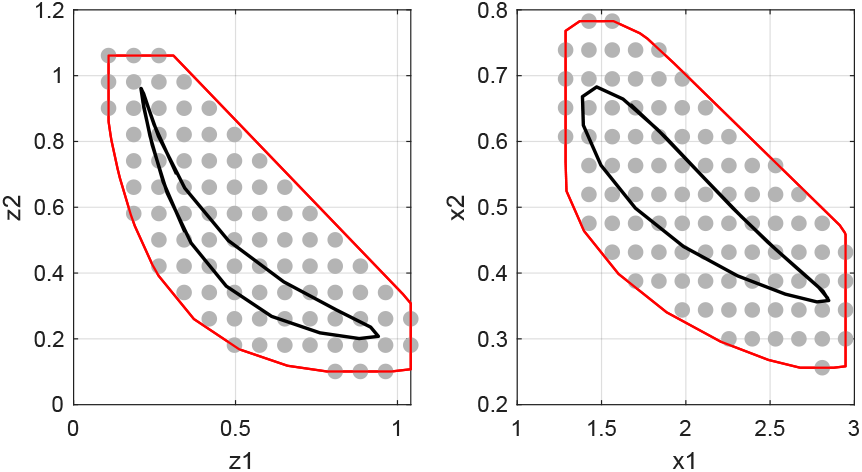
Region of feasibility of the LMI (8) for the closed loop (1), (3), (20). In contrast to our previous cases, the nonlinear binding between *x*_1_ and *x*_2_ and between *x*_1_ and *z*_1_, respectively, leads to a region of feasibility that constraints both *z* and *x* species. Dominance is certified by the feasibility of a set of LMIs built from the linearization at each shaded point.

At technical level, we did not address how to build regions of LMI feasibility that contain the desired attractor / trajectories. This is fairly straightforward in practice, starting from simulations and building sufficiently large regions that safely contain the trajectories of interest. However, the analysis can be made more rigorous by building (compact) regions of feasibility that are also forward invariant for the closed-loop dynamics. This additional property would entail the stronger results that every trajectory starting in the region must converge to some (simple) attractor within the region.

## 6. CONCLUSION

The results presented here show how to use tools from the dominance theory to study nonlinear systems in synthetic biology. We analyzed a particular nonlinear circuit architecture, the antithetic integral feedback system, which is shown to be able to encode homeostatic regulation (0-dominance) and robust periodic oscillations (2-dominance). For both cases, intuitive arguments based on the Nyquist criterion and root locus adapted to the linearized dynamics support parameter selection. Formal certificates are then provided by linear matrix inequalities. These certificates are inherently regional, in that they require the specification of a particular range of both state and parameter space. Remarkably, the approach allows us to make statements about robustness for oscillatory regimes in much the same way we use classical robust control to analyze the robustness of equilibria.

Overall, our paper support two important ideas. First, that the AIF circuit should be thought of as a core component in synthetic biology because of its capacity for diverse steady-state behaviors. From this perspective, we might think the AIF circuit as the biological equivalent of an op-amp, playing a central role in enabling monostable, multistable, and oscillatory circuits in synthetic biology. Second, that these behaviors structurally arise from nonlinearity and require control tools that go beyond the stability analysis of a single equilibrium. Dominance theory makes useful steps in this direction.

Our analysis is by no means comprehensive. We had made several simplifying assumptions and our analysis is limited to the specific set of parameters considered. However, we believe that the methodology described in this work presents progress towards a general approach to biological systems analysis. Dominance theory does not rely on the specific features of the nonlinearities, which makes it particularly well suited to biology, where models can be quite diverse and complex, yet the resulting dynamics are often surprisingly orderly and low-dimensional.

## ACKNOWLEDGEMENTS

The authors would like to acknowledge the support of Johan Paulsson in the Systems Biology department at Harvard Medical School and Rodolphe Sepulchre in the Department of Engineering at the University of Cambridge.

## Footnotes

1 For simplicity, all functions in the paper are assumed to be differentiable.

